# The dental calculus metabolome in modern and historic samples

**DOI:** 10.1101/136176

**Authors:** Irina M. Velsko, Katherine A. Overmyer, Camilla Speller, Matthew Collins, Louise Loe, Laurent A. F. Frantz, Juan Bautista Rodriguez Martinez, Eros Chaves, Lauren Klaus, Krithivasan Sankaranarayanan, Cecil M. Lewis, Joshua J. Coon, Greger Larson, Christina Warinner

## Abstract

**Introduction:** Dental calculus is a mineralized microbial dental plaque biofilm that forms throughout life by precipitation of salivary calcium salts. Successive cycles of dental plaque growth and calcification make it an unusually well-preserved, long-term record of host-microbial interaction in the archaeological record. Recent studies have confirmed the survival of authentic ancient DNA and proteins within historic and prehistoric dental calculus, making it a promising substrate for investigating oral microbiome evolution via direct measurement and comparison of modern and ancient specimens.

**Objective:** We present the first comprehensive characterization of the human dental calculus metabolome using a multi-platform approach.

**Methods:** Ultra performance liquid chromatography-tandem mass spectrometry (UPLC-MS/MS) quantified 285 metabolites in modern and historic (200 years old) dental calculus, including metabolites of drug and dietary origin. A subset of historic samples was additionally analyzed by high-resolution gas chromatography-MS (GC-MS) and UPLC-MS/MS for further characterization of polar metabolites and lipids, respectively. Metabolite profiles of modern and historic calculus were compared to identify patterns of persistence and loss.

**Results:** Dipeptides, free amino acids, free nucleotides, and carbohydrates substantially decrease in abundance and ubiquity in archaeological samples, with some exceptions. Lipids generally persist, and saturated and mono-unsaturated medium and long chain fatty acids appear to be well-preserved, while metabolic derivatives related to oxidation and chemical degradation are found at higher levels in archaeological dental calculus than fresh samples.

**Conclusions:** The results of this study indicate that certain metabolite classes have higher potential for recovery over long time scales and may serve as appropriate targets for oral microbiome evolutionary studies.

## 1. Introduction

Metabolites are small molecular weight molecules produced by a diverse range of enzymatic and chemical reactions, and include products derived from both endogenous and exogenous sources. Profiling metabolites in biological systems to define a metabolome is increasingly common, as it can provide insight into normal and perturbed metabolic processes and their relation to health and disease. Easily obtained human biofluids including serum (Psychogios et al. 2011), urine (Bouatra et al. 2013), and saliva (Dame et al. 2015) have been extensively profiled to define the range of metabolites that are produced in health, and how their levels fluctuate based on changes in activity (Daskalaki et al. 2015), diet (Lankinen et al. 2014), drug use (Fleet et al. 2008; Hahn et al. 1972), and disease progression (Yan et al. 2008). These studies have made it possible to search for specific metabolites that can act as biomarkers for a diverse range of disorders and diseases, including cardiovascular disease (Jensen et al. 2014), diabetes (Sysi-Aho et al. 2011), periodontal disease (Barnes et al. 2011; 2009), and cancer (Beger 2013).

Saliva has become an increasingly popular source for metabolite analysis because collection is simple, non-invasive, does not require training, and it is abundant and easy to resample and store (Dame et al. 2015). Saliva is composed mainly of water, but it also contains a wealth of molecules including mucins, proteins, carbohydrates, salts, and metabolites derived from serum, local cellular processes, diet, and oral microbes (Zhang et al. 2012). Because of the presence of serum-derived molecules, saliva has been used to search for biomarkers both of local diseases, such as periodontal disease (Barnes et al. 2011) and oral cancer (Yan et al. 2008), and also systemic diseases such as pancreatic and breast cancers (Sugimoto et al. 2010), and cardiovascular disease (Foley et al. 2012).

Metabolites from dental plaque, a microbial biofilm that develops on teeth, may provide novel information regarding host-microbiome interactions in health and disease. Dental plaque is likely to contain host saliva-and gingival crevicular fluid (GCF)-derived metabolites in addition to microbial metabolites, potentially providing a substrate for direct comparison of host and microbial activities. For reasons not well understood, dental plaque periodically and rapidly mineralizes to form dental calculus, a substance with concrete-like hardness that is immediately re-colonized by oral bacteria in a repetitive process (White 1991). Such rapid entombment has the potential to trap biomolecules from GCF and saliva as well as oral plaque and dietary and environmental debris (Warinner 2016; Warinner et al. 2015).

Although generally kept to low levels by professional dental hygiene regimens today, dental calculus was ubiquitous and relatively abundant in past human populations, as attested by dental calculus preserved within archaeological and paleontological collections spanning tens of thousands of years, and it is also found on the dentitions of some animal species (Warinner 2016; Warinner et al. 2015). Recent biomolecular investigations of ancient dental calculus have demonstrated that it contains exceptionally well preserved DNA and proteins from oral biofilm species, dietary components, and the host (Warinner, Hendy, et al. 2014; Warinner, Rodrigues, et al. 2014), as well as preserved plant microfossils (e.g., pollen, starch granules) and metabolic products (e.g, terpenoids) likely originating from dietary and craft activities (Blatt et al. 2011; Buckley et al. 2014; Hardy et al. 2012; Radini et al. 2016; Warinner, Rodrigues, et al. 2014). Such samples allow deep-time genetic and non-genetic molecular anthropology approaches to studying changes in human behavior, evolution of the oral biofilm and disease processes, and co-evolution of the oral microbiome and host, which are difficult to study using current *in vitro* and *in vivo* technologies alone. Gas-chromatography analyses of dental calculus from Neanderthals (Buckley et al. 2014; Radini et al. 2016), pre-agriculturalists (Hardy et al. 2016), and early agriculturalists (Hardy et al. 2012) have been used to infer the use of dietary and/or medicinal plants; however, to our knowledge, no broad-scale analysis to determine the potential range of metabolites trapped in dental calculus has been undertaken. Here we present an in-depth, shotgun metabolic analysis of a set of historic and modern dental calculus samples to assess the range of metabolites that can be extracted from calculus, and how well they persist and preserve over time. We validated our results and performed targeted metabolite searches in a subset of historic samples for a more thorough assessment of the potential preservation of metabolites of interest.

## 2. Materials and Methods

### 2.1 Calculus collection and preparation

Fresh dental calculus samples were obtained during routine dental cleaning at the University of Oklahoma Periodontology Clinic (n=1) in Oklahoma City, Oklahoma, USA and at a private dentistry practice (n=4) in Jaen, Spain (Table 1). Samples were collected by dental professionals using a dental scaler following standard calculus removal procedures. All samples were obtained under informed consent, and this research was reviewed and approved by the University of Oklahoma Health Sciences Center Institutional Review Board (IRB# 4543). Historic dental calculus (Figure S1a) was collected from 12 skeletons in the Radcliffe Infirmary Burial Ground collection, housed at Oxford Archaeology in Oxford, UK (Table 1). This cemetery was used from 1770-c.1855, and the skeletons are not personally identifiable. The oral health of each skeleton was recorded with reference to the presence or absence of caries and periodontal disease, with reference to (Hillson 1996; Ogden 2005). The sex and approximate age at death for each skeleton was estimated following standard osteological criteria (Brooks and Suchey 1990; Buckberry and Chamberlain 2002; Ferembach et al. 1980; Lovejoy et al. 1985; Phenice 1969; Schwartz 1996) and is presented in Table 1. Genetic sex was further confirmed through high-throughput sequencing (HTS) of DNA extracted from additional calculus fragments (described below) following previously described methods (Frantz et al. 2016; Skoglund et al. 2015; 2013); genetic sex determinations for all twelve samples were consistent with those made using osteological approaches. For details see Supplemental Methods.

**Table 1.** Summary of sample demographic and health information and analyses performed

| ID | Age<sup>a</sup> | Sex<sup>b</sup> | Smoker | Oral Health | Sampled Tooth | Analyses performed |
| --- | --- | --- | --- | --- | --- | --- |
| *Modern* |  |  |  |  |  |  |
| OK1010 | 40 | M | Former | + PD | - caries, + PD | UPLC-MS/MS |
| ES28 | 31 | F | Current | + caries, + GV | - caries, - PD | UPLC-MS/MS |
| ES29 | 30 | M | No | + caries, + GV | - caries, - PD | UPLC-MS/MS |
| ES20 | 23 | M | Current | + caries, + GV | - caries, - PD | UPLC-MS/MS |
| ES15 | 71 | M | Former | + caries, + PD | - caries, + PD | UPLC-MS/MS |
| *Historic* |  |  |  |  |  |  |
| CS06 | 36-45 | M | N/A | + caries, + PD | - caries, - PD | UPLC-MS/MS<sup>d</sup>; GC-MS |
| CS12 | 36-45 | M | N/A | + caries, + PD | - caries, - PD | UPLC-MS/MS<sup>d</sup>; GC-MS |
| CS18 | 36-45 | M | N/A | - caries, + PD | - caries, - PD | UPLC-MS/MS<sup>d</sup>; GC-MS |
| CS24 | 36-45 | F | N/A | + caries, + PD | + caries, + PD | UPLC-MS/MS<sup>d</sup>; GC-MS |
| CS46 | >45 | F | N/A | + caries, + PD | - caries, - PD | UPLC-MS/MS<sup>d</sup>; GC-MS |
| CS20 | 36-45 | M | N/A | + caries, + PD | - caries, - PD | UPLC-MS/MS |
| CS21 | 26-35 | M | N/A | - caries, - PD | - caries, - PD | UPLC-MS/MS |
| CS23 | 26-35 | F | N/A | + caries, + PD | - caries, - PD | UPLC-MS/MS |
| CS30 | 18-25 | F | N/A | + caries, - PD | - caries, - PD | UPLC-MS/MS |
| CS31 | 36-45 | F | N/A | + caries, + PD | - caries, - PD | UPLC-MS/MS |
| CS39 | 36-45 | M | N/A | - caries, + PD | - caries, - PD | UPLC-MS/MS |
| CS40 | >45 | F | N/A | - caries, - PD | - caries, - PD | UPLC-MS/MS |

<sup>a</sup>Age of skeletons used for historical calculus sampling is based on osteological indicators and is a range estimate.
<sup>b</sup>Historical skeleton sex estimate is based on both osteological indicators and DNA sequencing, and was in concurrence between the two methods. <sup>d</sup>UPLC-MS/MS was performed twice; the first at Metabolon, Inc. and the second at the University of Wisconsin-Madison. PD, periodontal disease; GV, gingivitis; N/A, not available; +, present; -, absent.

After collection, the fresh and historic dental calculus samples were stored frozen and transferred on dry ice to Metabolon, Inc. (Durham, NC) for sample processing and metabolite extraction and detection by UPLC-MS/MS. A subset of five historic dental calculus samples (CS6, CS12, CS18, CS24, and CS46) were additionally analyzed at the Departments of Chemistry and Biomolecular Chemistry at the University of Wisconsin (Madison, USA) to further characterize polar metabolites and lipids by high-resolution gas chromatography (GC)-MS and UPLC-MS/MS, respectively (Table 1).

### 2.2 Genetic Authentication of a Preserved Oral Microbiome in Historic Samples

DNA extracted from a separate fragment of each piece of historic calculus was used to assess microbial community composition. DNA was extracted as previously described, but omitting phenol-chloroform steps (Warinner, Rodrigues, et al. 2014), and Illumina shotgun sequenced (for details see Supplemental Methods). The 16S rRNA gene-identified reads were then used to assess microbial community composition at the genus level by closed-reference OTU-picking against the GreenGenes v. 13.8 database using UCLUST (Edgar 2010) in QIIME v. 1.9 (Caporaso et al. 2010). The Bayesian analysis-based program SourceTracker (Knights et al. 2011) was used to estimate the source composition of the microbial community identified by QIIME. Human reads were identified by mapping to GRCh38.p10 (GCF_000001405.36) using bwa (Meyer et al. 2012) with the flags aln -l 16500 -o 2 -n 0.01, duplicate reads were removed, and reads mapping to X and Y chromosomes were analyzed for genetic sex determination as described above.

### 2.3 Mass spectrometry and data processing for UPLC-MS/MS

Samples (∼20 mg) were decalcified in 0.5M EDTA, centrifuged to pellet debris, and supernatant prepared using the automated MicroLab STAR® system from Hamilton Company. Samples were cleaned and divided into five fractions: two for analysis by two separate reverse phase (RP)/UPLC-MS/MS methods with positive ion mode electrospray ionization (ESI), one for analysis by RP/UPLC-MS/MS with negative ion mode ESI, one for analysis by HILIC/UPLC-MS/MS with negative ion mode ESI, and one sample was reserved for backup. All methods utilized a Waters ACQUITY ultra-performance liquid chromatography (UPLC) and a Thermo Scientific Q-Exactive high resolution/accurate mass spectrometer interfaced with a heated electrospray ionization (HESI-II) source and Orbitrap mass analyzer operated at 35,000 mass resolution. For details see Supplemental Methods. Raw data was extracted, peak-identified and QC processed using Metabolon’s hardware and software. Compounds were identified by comparison to library entries of purified standards or recurrent unknown entities. Peaks were quantified using area-under-the-curve. Each compound was corrected in run-day blocks by registering the medians to equal one (1.00) and normalizing each data point proportionately.

### 2.7 Further characterization of historic calculus by GC-MS and UPLC-MS/MS

Five historic dental calculus samples were selected to further investigate polar metabolites and lipids in historic dental calculus samples (Table 1). Following sample pulverization, 15 mg was decalcified with 100 uL of 4% Formic acid. Samples were incubated at 4°C with occasional shaking for 12 days. Next, 75 uL of 1 M ammonium hydroxide was added, then samples were extracted with 350 uL MeOH + 350 uL Acetonitrile (final 2:2:1 Methanol:Acetonitrile:Water). Extract was split for polar metabolite (GC-MS) and lipid (LC-MS) analysis and dried down by vacuum centrifugation. For GC-MS analysis molecules were analyzed with positive electron-impact (EI)-Orbitrap full scan of 50-650 m/z range. Lipid LC-MS analysis was performed on a Water’s Acquity UPLC CSH C18 Column (2.1 mm x 100 mm) with a 5 mm VanGuard Pre-Column Mobile coupled to a Q Exactive Focus. Raw files were analyzed using Thermo Scientific’s Tracefinder 4.0 deconvolution plugin and unknown screening quantification tool (GC-MS), or the Thermo Compound Discoverer^TM^ & 2.0 application with peak detection, retention time alignment, and gap filling (UPLC-MS/MS). Only peaks 10-fold greater than solvent blanks were included in the later analysis. For details see Supplemental Methods.

### 2.8 Data analysis

Mass normalized data was used for all downstream analyses. First, overall metabolome composition was summarized at the super-pathway, sub-pathway, and metabolite levels, and identified metabolites were cross-referenced against public databases to obtain KEGG compound identifiers and Human Metabolome Database (HMDB) IDs. Next, metabolites found to be ubiquitously present in modern samples and ubiquitously absent in historic calculus were compared. Here ubiquity among the five modern samples was applied as a measure to identify highly prevalent (potentially core) dental calculus metabolites; complete absence of these metabolites among all twelve historic samples was used to identify metabolite candidates that may be particularly prone to loss or that are unstable and susceptible to degradation through time-dependent taphonomic processes. Following this analysis, differential representation of metabolites between historic and modern samples was determined using the program Statistical Analysis of Metagenomic Profiles (STAMP) (Parks and Beiko 2010; Parks et al. 2014), first including metabolites detected in both historic and modern samples, and then again using only those metabolites that were universally detected in all seventeen calculus samples. Metabolite profiles of historic and modern samples were compared using 2 group analysis of the average quantity of each metabolite and analyzed by White’s non-parametric two-sided t-test with bootstrapping to determine the difference between proportion (DP) with cut-off 95% and Storey’s FDR. Differential abundance was determined in hierarchical categorization of super-pathway, sub-pathway, and individual metabolite, for the mean relative proportion of metabolite(s) at each level by two-sided Fisher’s exact test using the Newcomb-Wilson DP with cut-off 95% and Storey’s FDR. For both analyses corrected p-values (q-values) of ≤ 0.05 together with an effect size ≥1 were considered significant. Pathway maps were created using iPATH2 (Yamada et al. 2011) for metabolites with KEGG compound identifiers. Partial Least Squares Discriminant Analysis (PLS-DA) was performed using the R package mixOmics (Rohart et al. 2017) in default settings.

## 3 Results

### 3.1 Authentication of historic calculus

Archaeological specimens are subject to environmental degradation and contamination, and thus it is necessary to confirm of the source (e.g., endogenous microbiome vs. exogenous environmental microbes) of biomolecules detected in ancient samples. QIIME and SourceTracker analyses confirmed excellent biomolecular preservation of an *in situ* oral microbial community within the historic dental calculus samples, and16S rRNA gene sequences closely matched those expected for dental plaque communities, with minimal contamination from exogenous sources such as soil and skin (Figure S1b). The high proportion of microbes of “unknown” source in several historic dental calculus samples is observed in modern calculus samples (Ziesemer et al. 2015) (Figure 1b ‘Modern’), and is a result of mismatched source samples. Several poorly taxonomically characterized oral taxa such as *Methanobrevibacter* and *Tissierellaceae* are highly abundant in mature dental calculus biofilms but are infrequently detected in healthy oral plaque early biofilms such as the Human Microbiome Project cohort we used as source samples. Therefore these genera cannot be confidently assigned to an oral plaque source, and are instead attributed to an unknown source.

**Fig. 1.**
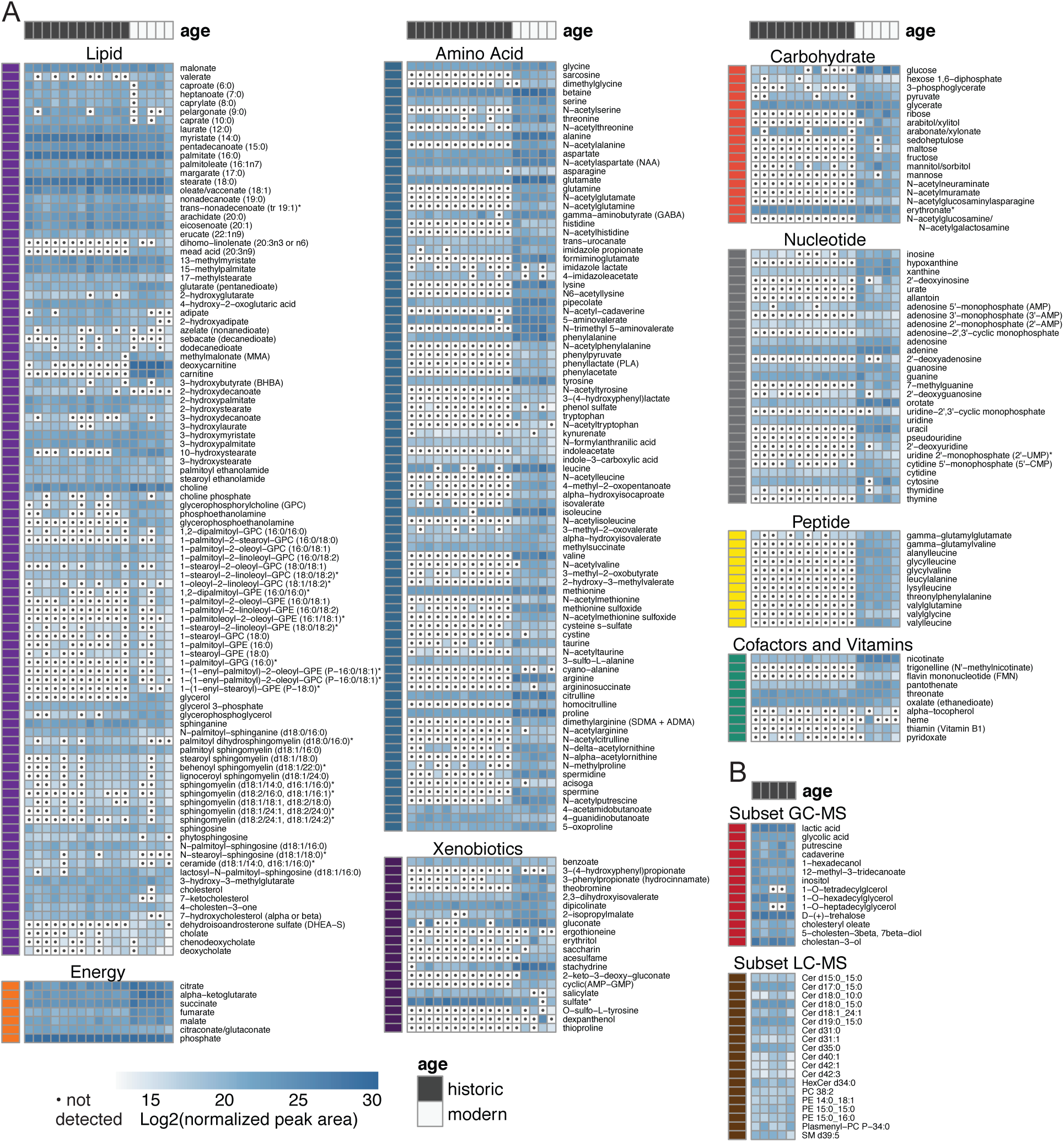
Heat map summary of metabolites observed in modern and historic dental. Metabolites were quantified by area under the curve and normalized to mass of sample extracted. **a** UPLC-MS/MS-detected metabolites. Samples were hierarchically clustered, and log2 transformed values are presented above, grouped by super-pathway. **b** Metabolites detected by GC-MS and LC-MS in historic calculus. Non-filled cells containing a dot indicate the compound was not detected.

### 3.2 Metabolic pathway coverage in dental calculus

A total of 285 metabolites were identified by UPLC-MS/MS in the seventeen dental calculus samples, and these were categorized as members of one of eight super-groups: *Amino acids, Carbohydrates, Cofactors and vitamins, Energy, Lipids, Nucleotides, Peptides*, and *Xenobiotics* (Figure 1a; Supplemental Table S1), which were further classified into 69 sub-categories. More than half of the metabolites (n=185) were detected in both historic and modern calculus samples, demonstrating that metabolites can be recovered from historic dental calculus, while 99 metabolites were detected only in modern samples and 1 was detected only in historic samples. One hundred ninety-nine metabolites were detected in all 5 modern samples, 97 were detected in all 12 historic samples, and 85 were detected in all 17 calculus samples (Table S1). A smaller subset of historic calculus was analyzed by GC-MS and LC-MS at the University of Wisconsin-Madison; this enabled the identification of 15 additional metabolites and 40 additional lipids, respectively (Figure 1b; Table S2). For metabolites that were quantified in the main analysis and in the smaller subset, we see comparable quantitation (positive correlation, R^2^ = 0.7, Supplemental Figure S2). UPLC-MS/MS-identified metabolites with KEGG compound identifiers (n=207) were located on a general metabolic pathway map (Figure S3) and a map of biosynthesis of secondary metabolites (Figure S4).

### 3.3 Comparison of dental calculus and saliva metabolomes

To determine the degree of overlap between metabolites in dental calculus and metabolites in saliva, we compared our results to the saliva metabolome. We downloaded a list of all metabolites reported in saliva as catalogued in the Human Metabolome Database (Wishart et al. 2013) version 3.6 (hmdb.ca) as of February 2017 to use as the known saliva metabolome. This list contained 1233 metabolites of endogenous and exogenous origin, spanning the full range of super-pathways detected in calculus samples. Just over half, 159, of the 285 metabolites detected in calculus (55.7%) were previously included in HMDB’s saliva metabolome (Table S1), while these 159 metabolites make up just 12.9% of the total metabolites detected in saliva. Of the remaining 107 metabolites detected in calculus, 84 have been detected in blood, urine and/or cerebrospinal fluid, and 23 have no HMDB identifier. At least one metabolite in each of the sub-pathways represented in calculus is not included in the HMDB saliva metabolome.

### 3.4 Metabolite preservation patterns

Among metabolites that are likely endogenous (host or oral microbiome) in origin (i.e., not xenobiotics), metabolite persistence differs by super-pathway in historic samples (Figure 1). Overall, historic samples had lower representation in metabolites categorized as *Amino acids*, *Vitamins and cofactors*, *Carbohydrates*, *Nucleotides*, and *Peptides* (Figure 2a). In contrast, *Lipids* and *Energy* metabolites were generally observed in both historic and modern calculus (Figure 2a). Additionally, at a finer scale, certain chemical configurations appear to be lost through time, for example N-acetylation, amino acids with positively-charged R-groups, and phenyl rings one carbon away from an oxygen. These data suggested a preservation bias that could be due to either chemical stability or compound solubility, although it is a possibility that low sample amounts limit the detection of certain metabolites.

**Fig. 2.**
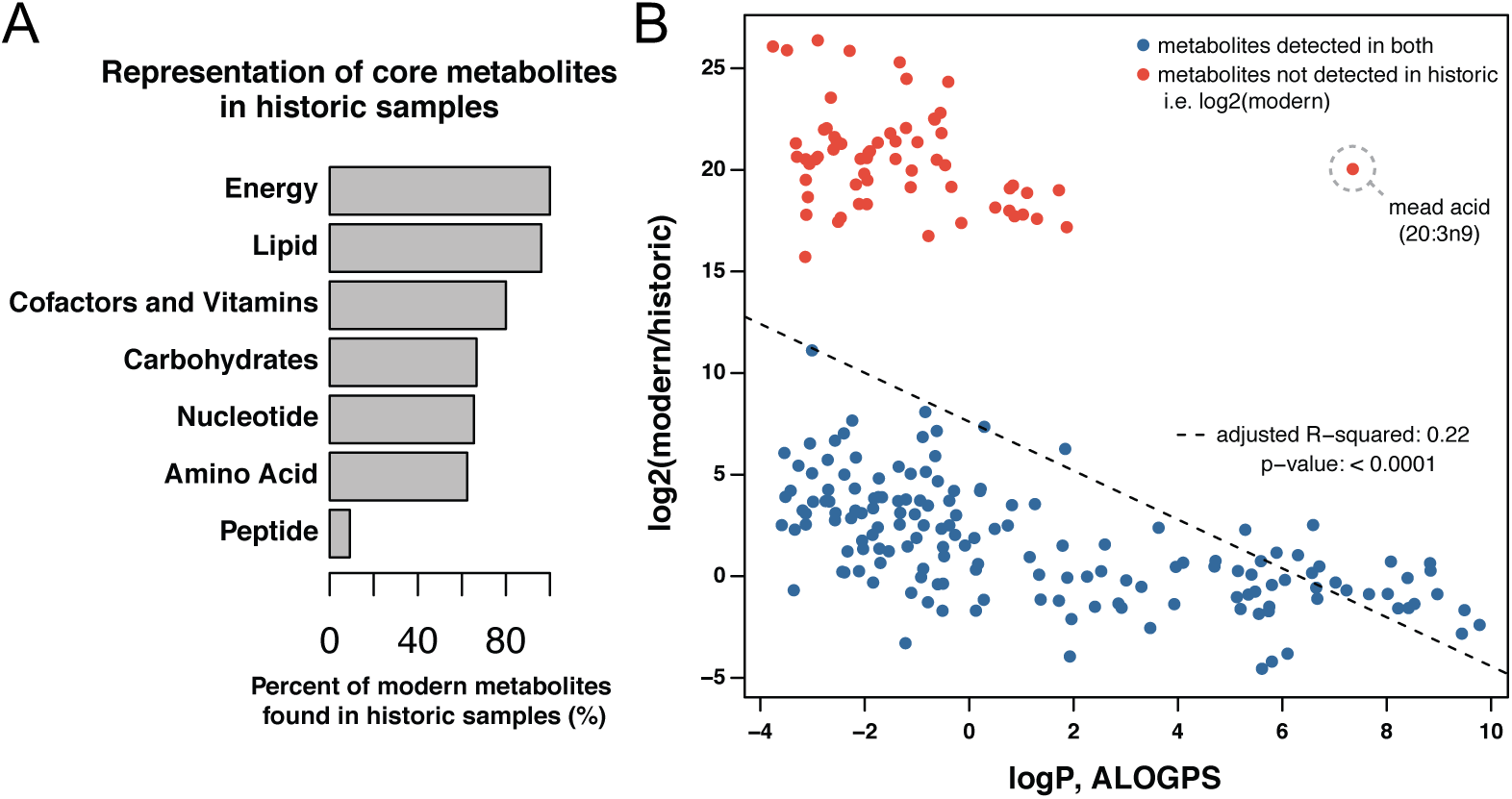
Compound preservation is correlated with aqueous solubility. **a** Percent of high ubiquity metabolites in modern calculus that were also recovered in at least one historic calculus sample. *Peptides* exhibit the poorest representation in historic dental calculus, with only 9% of the peptides observed to be present in all five modern calculus samples also detected in any historic sample. By contrast, *Lipids* and *Energy* (TCA cycle) super-pathways exhibit high representation in historic calculus, with >96% of compounds found in all five modern samples also recovered from historic dental calculus. *Xenobiotics*, which largely comprise dietary and pharmaceutical compounds, are not shown. **b** The log2 fold-change (modern/historic) of metabolite abundance vs. the 1-octanol vs. water partition coefficient (logP), estimated with the ALOGPS tool. In the cases where metabolites were only detected in modern calculus, the log2 values (not fold-change) were plotted relative to logP. The fitted linear model showed a significant effect (p<0.0001) of logP on metabolite fold-change with and adjusted R^2^ of 0.22. The outlier from this trend was mead acid (20:3n9).

To test the hypothesis that preservation is linked to aqueous solubility, we compared the differential abundance of metabolites between modern and historic calculus to the predicted hydrophobicity of the compounds. The predicted hydrophobicity was extracted from the HMDB, which sources the ALOGPS predicted ratio of compound partitioning between 1-octanol and water (logP) (Tetko and Bruneau 2004). When plotting fold-change abundance (modern/historic) to logP, we observe a significant (p<0.001) negative correlation (Figure 2b), suggesting that molecules that are more abundant in modern calculus have lower organic solubility and higher aqueous solubility. This result is consistent with our hypothesis that metabolite preservation is in part due to aqueous solubility. The exception to our hypothesis, and the outlier in Figure 2b, are poly-unsaturated fatty acids (PUFAs), which have high logP but low preservation. Two PUFAs, mead acid (20:3n9) and dihomo-linolenate (20:3n3 or n6), were identified in all modern dental calculus samples but were not observed in historic samples. The loss of these PUFAs in historic calculus may be partially explained by the decreased oxidative stability of fatty acids with decreasing saturation (Cosgrove et al. 1987; Rustan and Drevon 2005).

As yet there are no detailed assessments of metabolite degradation in archaeological dental calculus; however, analyses of protein damage patterns in human dental calculus and mammoth bone (Cappellini et al. 2012) can be used to draw comparisons with damage patterns in calculus metabolites. Warinner, *et al*. (Warinner, Rodrigues, et al. 2014) found that the most common protein post-translational modification products in ancient dental calculus are deamidation of asparagine, deamidation of glutamine, oxidation of methionine, and conversion of N-terminal glutamine to pyroglutamate. Asparagine was detected in all five modern samples and eleven historic samples, while the deamidation product aspartate was detected in all seventeen samples, and in higher quantity. Glutamine was detected in all modern samples but not in historic samples, and its deamination products glutamate and 5-oxoproline were detected in higher concentration in all 17 samples, although at much higher concentration in modern than historic samples. Oxidation, which is widely documented in the degradation of oil paintings (Oakley et al. 2015) and food spoilage (Velasco and Dobarganes 2002) also occurs in dental calculus. For example, the ratio of cholesterol and its oxidation product 7-ketocholesterol was reversed between modern and historic calculus, and kynurenin, an oxidation product of tryptophan that is known to accumulate in archaeological bone over time (Cappellini et al. 2012), was detected in historic calculus while tryptophan was absent. In contrast, methionine was detected in all seventeen samples, and was much more abundant in modern samples, while the oxidation product methionine sulfoxide was detected in all five modern samples and in only two historic samples, in all cases at lower concentrations. Free methionine sulfoxide may be unstable and subject to further rapid breakdown.

### 3.5 Lipid 2-hydroxylation as an indicator of calculus age

Four 2-hydroxylated lipids were detected in all calculus samples—2-hydroxyadipate, 2-hydroxystearate, 2-hydroxypalmitate, and 2-hydroxyglutarate—and the first three are more abundant in ancient than modern samples. The only metabolite detected in historic but not modern samples was 2-hydroxydecanoate (detected in 10 of the 12 historic samples), while the parent molecule decanoate was not detected in any of our samples. In contrast, 3-hydroxylated and 2,3-hydroxylated lipids are more abundant in modern than historic samples. The increased presence of 2-hydroxylated lipids in ancient samples suggests that this modification may increase over time. The difference in patterns of lipid hydroxylation between ancient and modern calculus suggests that in some cases 3-hydroxylation may switch to 2-hydroxylation.

### 3.6 Differentially abundant metabolites in ancient and modern calculus

To further define the differences in metabolic functions preserved through time, we compared the metabolites present in both modern and ancient calculus samples at the super-pathway, sub-pathway, and individual metabolite levels using STAMP. Principal components analysis demonstrated distinct separation of modern and historic samples (Figure 3a) with tight clustering of historic samples along PC1 and PC2, while modern samples were more distributed, suggesting that loss of metabolites through time results in a more uniform metabolite profile between samples than may have originally existed. Comparing the mean proportions of individual metabolites detected in at least one modern and one historic sample, 161 were significantly more abundant (q ≤ 0.05) in one sample set, yet only 21 additionally had an effect size of ≥1.0 (Figure 3b, Table S3 bold metabolites). When considering only the metabolites that were universally present in all five modern and all twelve historic samples (Table S3, superscript ‘c’), a slightly different set of metabolites and metabolic pathways are differentially abundant. Historic and modern samples still separate distinctly in PCA (Figure S5a), but many more metabolites have a significant difference in mean proportions (Figure S5b).

**Fig. 3.**
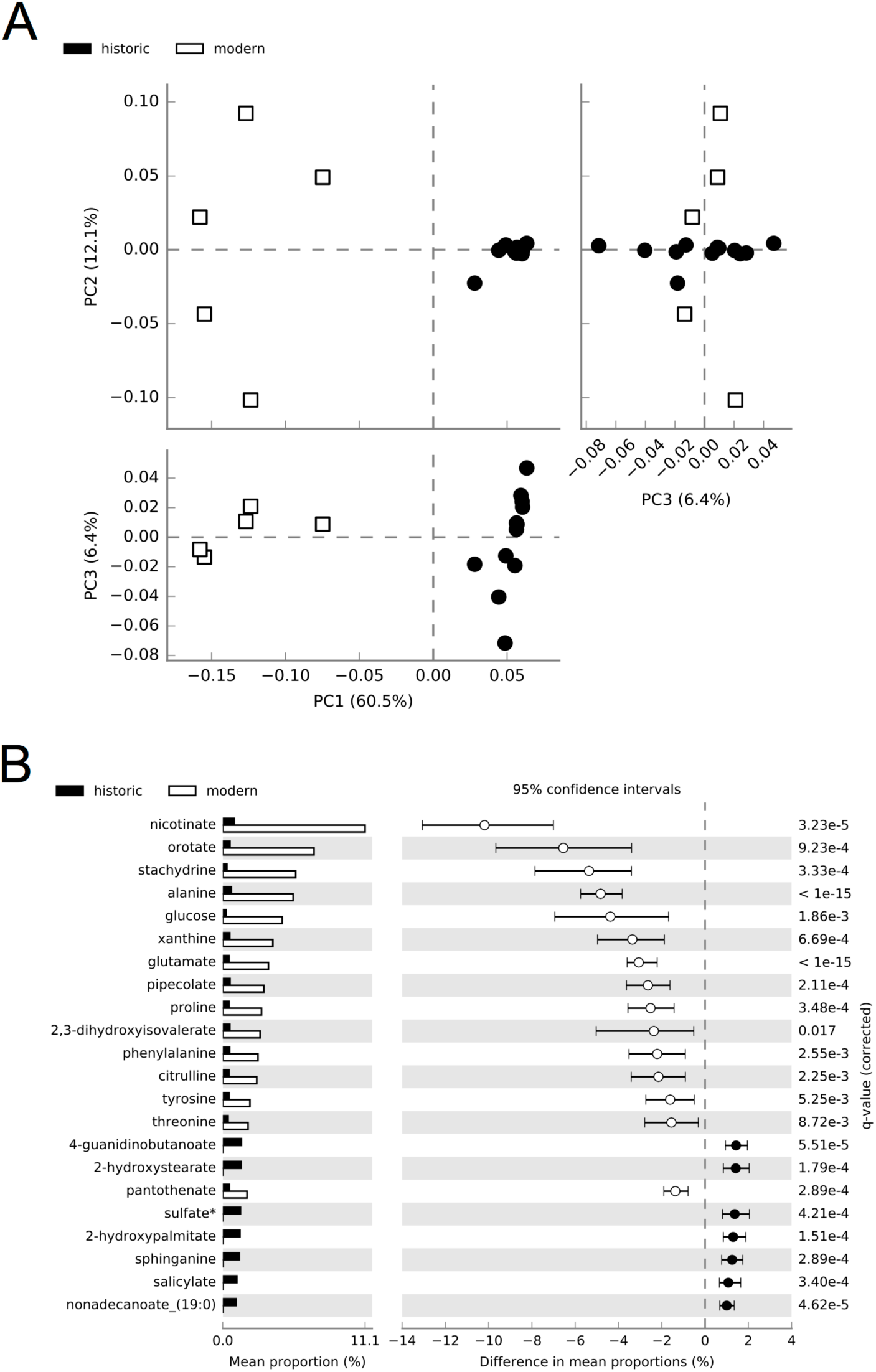
Differences exist in mean proportions of metabolites detected in at least one historic and one modern dental calculus sample. **a** Principal components analysis distinctly separates modern and historic calculus samples. **b** Metabolites with significant differences (q ≤ 0.05, effect size of ≥1.0) in mean proportions between historic and modern calculus.

We then examined differences in proportion of super-pathways, sub-pathways, and individual metabolites present in at least one modern and historic sample to better understand the patterns of preservation and loss through time. The proportion of *Lipids* was significantly higher in ancient than modern calculus samples, suggesting that non-polar, chemically inert molecules are particularly stable through time (Figure 4a). On the other hand, the proportion of *Amino acids*, *Carbohydrates*, *Cofactors and vitamins*, and *Xenobiotic*s were significantly higher in modern calculus (Figure 4a), demonstrating substantial loss and/or degradation of metabolites in these super-pathways over time. The super-pathway *Peptides* was excluded from analysis using this method because its near total absence in historic samples resulted in very few possible comparisons.

**Fig. 4.**
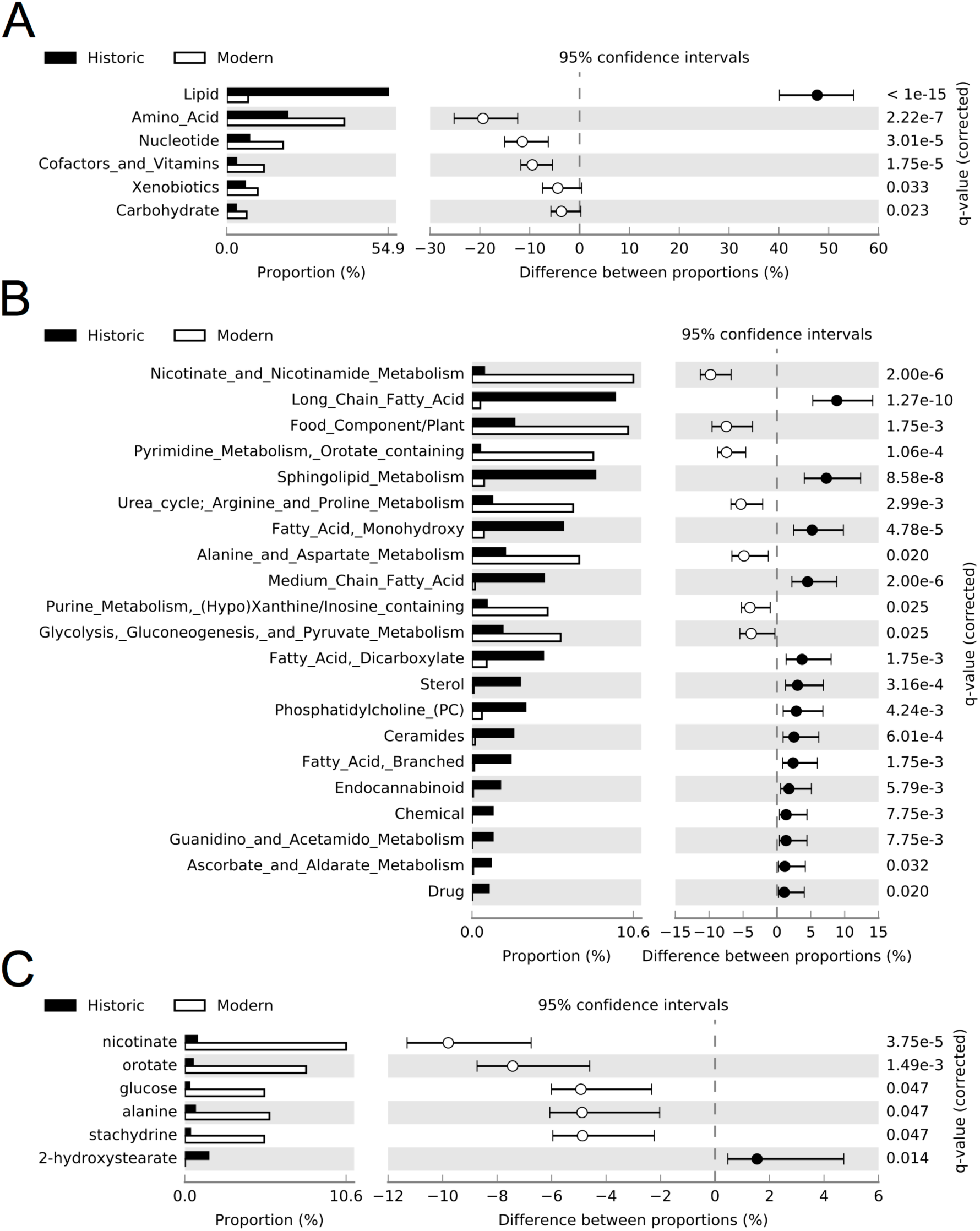
Differences exist in proportions of super-pathways, sub-pathways, and metabolites represented in at least one historic and one modern dental calculus sample. Significantly different (q ≤ 0.05, effect size of ≥1.0) proportions of **a** Super-pathways, **b** Sub-pathways, and **c** metabolites. Individual metabolites between historic and modern samples.

As expected from the super-pathway differential abundances, many of the sub-pathways with greater proportional representation in historic calculus were related to lipid metabolism (Figure 4b). The sterols include cholesterol and its oxidation products 4-cholesten-3-one, 7-hydroxycholesterol (alpha or beta), and 7-ketocholesterol, which is consistent with the expectation that increased oxidation will occur over time. *Guanidino and acetamido metabolism* comprised a greater proportion of historic sample metabolites due to overrepresentation of 4-guanidinobutanoate, a product of arginine and putrescine metabolism. Several of the sub-pathways with greater proportional representation in modern samples can be explained by a single metabolite, which manifests at the metabolite level (Figure 4b), and these include *Nicotinate and nicotinamide metabolism*, *Pyrimidine metabolism*, and *Food component/plant*. A single metabolite each comprises the *Chemical* and *Drug* sub-pathways, sulfate and salicylate, respectively. With respect to the latter, salicylate is abundant in modern pharmaceuticals, but may have been consumed medicinally in the form willow bark tea by the historical population, especially given the fact that they were buried in a hospital-associated cemetery. Salicylates are also naturally found in a variety of fruits, vegetables, herbs, and spices; however at levels far below the therapeutic doses typical of modern pharmaceuticals (Castillo-García et al. 2015).

In contrast to the large number of sub-pathways representing a significantly higher proportion of metabolites between historic and modern calculus, only 6 individual metabolites were significantly differentially represented between historic and modern calculus (Figure 4c). Modern calculus had significantly higher proportions of nicotinate, orotate, stachydrine, alanine, and glucose (Figure 4c). Both nicotinate and orotate may be taken as a dietary supplement, while stachydrine (proline betaine), is a plant metabolite that is not metabolized by mammals (Lever et al. 1994), but is common in citrus fruits and orange juice (Atkinson et al. 2007). Low abundance of stachydrine in historic calculus may relate to dietary differences between these modern and historic populations, or to differential preservation. Alanine, the smallest amino acid, and glucose may be lost through high solubility. The lipid 2-hydroxystearate was the only metabolite of significantly greater proportion in historic samples, even though several 2-hydroxylated lipids had greater relative abundance in historic samples.

In addition to endogenous metabolites, several xenobiotics found to be present only in modern calculus samples are from food and pharmacologic agents introduced to or popularized in European populations in the 20^th^ century, including the artificial sweeteners acesulfame, saccharin, and arabitol/xylitol. Additionally, theobromine, an alkyloid present in coffee, tea, and chocolate, was also only detected in modern calculus, suggesting that consumption of these products by the historic population was absent or low, despite their increasing availability in Europe in the 1800s, or that this metabolite poorly preserves over time.

### 3.7 Potential for maintenance of biological signatures in calculus metabolite profiles

Unlike saliva, dental calculus does not represent a snapshot of a specific time and specific metabolic state, but rather it represents a life history in which specific profiles may be diluted out by fluctuating metabolic processes throughout an individual’s life, loss of unstable metabolites over time, and random chance with respect to the entrapment of xenobiotics and dietary compounds. However, distinct metabolic signatures related to biological variables such as sex (Takeda et al. 2009), oral health status (Barnes et al. 2009; 2011), and oral biofilm microbial composition (Takahashi et al. 2010) have been reported in saliva, GCF, and oral plaque samples, all of which are likely to contribute to the metabolite profile in dental calculus. Therefore, we assessed differences in metabolic profiles between calculus from different age groups, sex, and oral health status by partial least squares discriminant analysis (PLS-DA), and, further, looked for metabolites that could be specifically attributed to bacterial activity.

No differences were found in the metabolite profile between age groups, between males and females, between samples from caries-affected and caries-free dentitions, or between samples from periodontal disease-affected and non-affected individuals when considering metabolites detected in at least one historic and one modern calculus sample (Figure 5a), or metabolites universally present in all 17 samples (Figure S7a). However, there was a distinct separation of modern and historic samples in each comparison so we repeated PLS-DA using only historic sample data. PLS-DA separated historic samples based on sex, age, caries status and periodontal disease status when using metabolites detected in at least one historic sample (Figure 5b) and when using only metabolites present in all 12 historic samples (Figure S7b). Applying PSL-DA to universally detected metabolites in only the *Lipid* and *Energy* classes, (the best-preserved classes in historic samples, Figure 1a) separated samples based on time period rather than biological category (Figure S7c). These results suggest that biological categories in modern and historic calculus samples cannot be directly compared, yet patterns of biological differences are maintained through time.

**Fig. 5.**
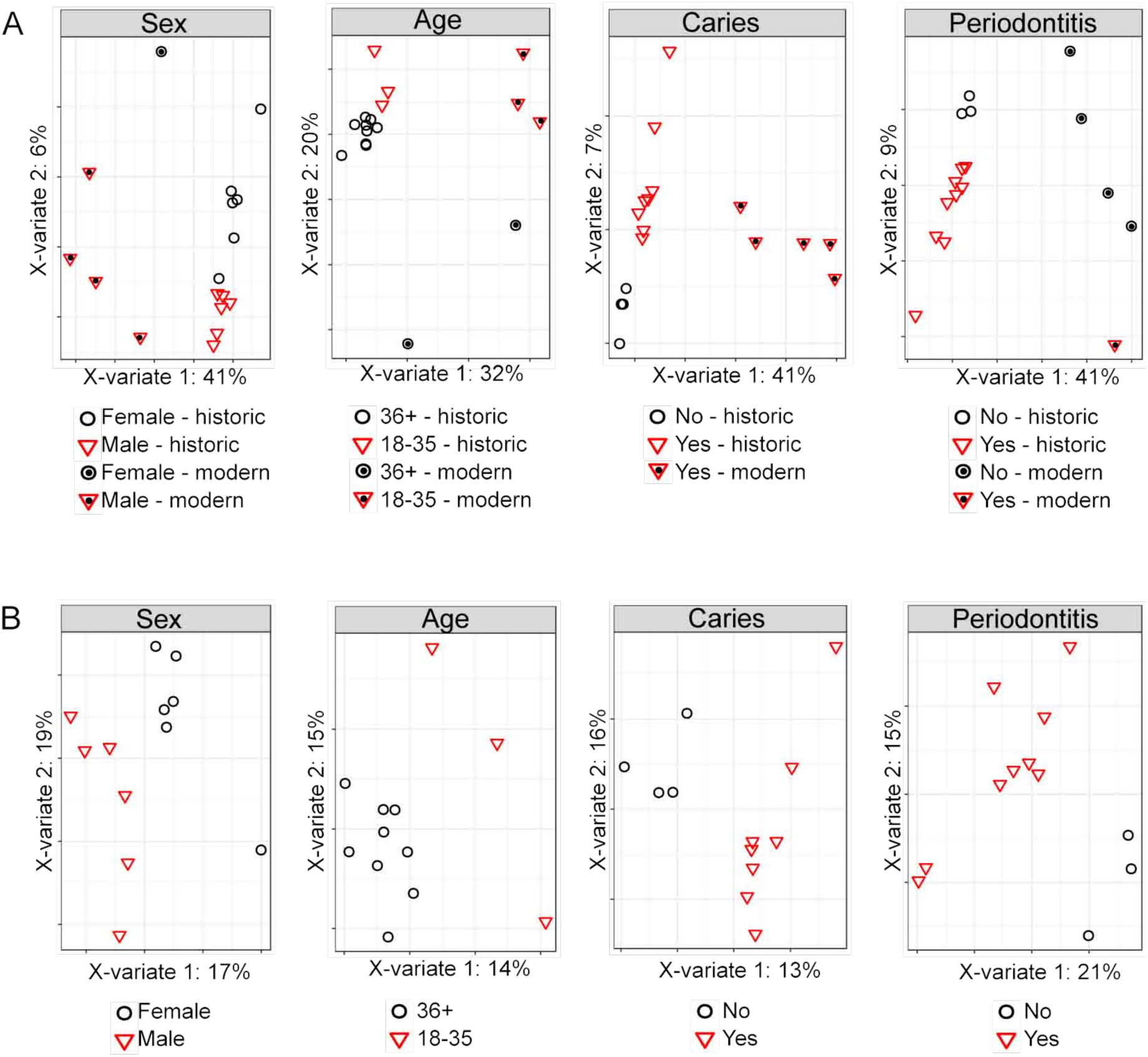
Partial least squares discriminant analysis of metabolites detected at least one modern and one historic calculus sample. **a** Calculus samples cluster based on time period rather than biological category (sex, age, caries status and periodontal disease status) when including all metabolites detected in at least one modern and one historic sample. **b** Historic calculus samples cluster based on biological category (sex, age, caries status and periodontal disease status) when including all metabolites detected in at least one historic sample.

Similarly, no metabolites could be specifically attributed to bacterial processes, but several metabolites, including isovalerate, valerate, lactate, cadaverine and putrescene, are known metabolic products of oral bacteria (Scully et al. 1997; Takahashi 2015).

Dipicolinate, which was detected in all modern and all historic samples, is the major component of bacterial spore capsules, and may indicate that bacterial endospore development occurs in mature plaque biofilms, or during plaque mineralization. Sulfate is abundant in GCF due to break-down of sulfur-containing amino acids by the oral biofilm, and is produced during anaerobic methanogenesis in oral plaque, however, we observed no correlation between the relative abundance of the oral methanogen *Methanobrevibacter* and sulfate levels (Figure S8) in our samples, and the very high abundance of sulfate in historic relative to modern calculus suggests an exogenous source. Oxford topsoil sulfur concentrations where the cemetery was located are in the 70^th^ percentile across England and Wales (Rawlins et al. 2012), and sulfates in the soil and ground water of the cemetery may leach into the calculus by the same processes through which highly water-soluble metabolites are leached out of calculus.

## 4 Discussion

Our results derived from the non-targeted assessment of metabolites present in dental calculus from both modern and historic samples demonstrate the significant potential of calculus as a material for metabolomics and lipidomic studies. The wide range of metabolic categories covered (amino acids, carbohydrates, cofactors and vitamins, energy, lipids, nucleic acids, peptides, xenobiotics), and the variety of sources of metabolites (host, microbial, diet) are on par with those reported in saliva (Barnes et al. 2011) and GCF (Barnes et al. 2009) using similar metabolite detection platforms. Similar to the study defining the saliva metabolome (Dame et al. 2015), we found using multiple metabolomics platforms (5 different methods by Metabolon, Inc. and 2 by UW-Madison) increased the diversity of compounds we detected. However, unlike Dame *et al* (2015), we found more metabolites by LC than GC methodologies, which was likely due to the low-abundance of molecules with higher aqueous solubility in the historic calculus that were analyzed by GC-MS. Although we found lower representation of aqueous-soluble molecules, in general calculus preserves a wide variety of molecules from the oral cavity and could be useful proxy for oral biofluids in archaeological samples. Calculus also provides an opportunity to co-investigate host and microbial activity, which is increasingly recognized as important to understanding cellular physiology and disease pathology (Takahashi 2015).

Saliva has been shown to preserve an individualized metabolic signature throughout daily routine (Wallner-Liebmann et al. 2016) and dental calculus, which contains salivary components, has the potential to preserve aspects of individual profiles over longer periods of time. While only five modern calculus samples were included in this study, PCA analysis revealed substantial metabolite profile diversity within these samples (Figure 2a). It therefore appears that modern calculus may preserve individual phenotypes, although to investigate this, studies with larger sample sizes are needed to further assess this potential. Historic samples, in contrast, cluster much more tightly in the PCA, suggesting that individual phenotypes may be lost through metabolite degradation and loss over time. However, we were able to distinguish metabolic profile differences in historic samples between sex, age, and oral health status of the individuals by PLS-DA, demonstrating maintenance of individual profiles despite metabolite loss. It may then be possible to investigate differences in specific metabolite profile in historic samples, which could contribute to our understanding of disease demographics and evolution.

Relatively little is known about the process of age-related protein degradation in archaeological samples, yet other historic samples provide some insight. Asparagine readily deamidates via cyclization to succinimidyl within chain; however this mechanism is unavailable to the free amino acid. Under experimental heating, asparagine undergoes rapid hydrolysis (Crisp et al. 2013), and it is therefore probable that the free asparagine seen in the historic samples is derived from hydrolysis of peptides. Free glutamine and glutamic acid can undergo cyclisation to pyroglutamic acid even at low temperatures (Nagana Gowda et al. 2015). However, although pyroglutamate (pGlu; 5-oxoproline) is present at higher levels in the historic samples, it is too low to account for all the loss of all glutamic acid. This consistent pattern suggests that there is a contributing pool of degrading proteins, generating free amino acids which are undergoing modification, either in chain (e.g., asparagine deamidation) or once hydrolysed to terminal positions or as free amino acids (e.g., pyroglutamate). The majority of these low molecular weight, high solubility products are then likely lost from the calculus matrix. It is possible that some are so entrapped within the crystal matrix that they may persist as free amino acids and pyroglutamate, and this could be assessed by monitoring the level of racemization (Crisp et al. 2013).

Lipids, particularly unmodified, saturated classes, are some of the best-preserved metabolites in historic calculus, and appear to be particularly stable over time. Therefore, lipid analyses may be a promising focus for historic calculus studies. Although not a common focus in salivary or oral biofilm metabolomics studies, lipids are a versatile class of molecule with a broad range of physiological properties and actions. They play roles in local (intracellular) (Nishizuka 1995) and long-distance (hormone) cell signaling (Xavier et al. 2016), have both pro-and anti-inflammatory properties (Bennett and Gilroy 2016), and are the major components of cell membranes, where their composition influences cell membrane function (Zalba and Hagen 2017). Bacterial membrane lipid content varies by species (López-Lara and Geiger 2016), and may indicate bacterial physiological status (Darveau et al. 2004), while pathogenesis of the periodontal disease-associated oral bacteria *Porphyromonas gingivalis* is influenced by host cell membrane lipid composition (Wang and Hajishengallis 2008). The role of lipid mediators in the initiation and resolution of periodontal disease inflammation is currently under investigation (Bartold and Van Dyke 2013), and the wealth of lipids detected in calculus may be valuable to studies of both host and microbial pathophysiology.

Although we were unable to specifically identify bacterial contributions to the calculus metabolome, there are some metabolites suggestive of mature oral biofilm activity. Dipicolinate, which is the major capsule component of bacterial endospores, is a highly stable molecule, as evidenced by the long-term viability of endospores (Yung et al. 2007). To our knowledge, the inferred presence of endospores in calculus is a novel finding, as we were unable to find any references to the presence of bacterial endospores in oral plaque or dental calculus. Members of several Gram-positive genera that reside in the mouth have close relatives known to form spores, including *Actinomyces* (Gao and Gupta 2012), and *Filifactor* (Vos et al. 2011), and since many oral bacteria have not yet been genetically or physiologically characterized, it is possible that several oral species do have the ability to form spores. Calcification of the biofilm may induce a stress response in these species that initiates endospore formation, which would explain the abundance of dipicolinate in dental calculus.

Additionally, studies aiming to characterize salivary biomarkers of periodontal disease have identified several pathways with an apparent bacterial source that contain promising metabolite candidates for disease biomarkers (Kuboniwa et al. 2016; Sakanaka et al. 2017), and we have identified several of these in our calculus. Phenylalanine, succinate, hydrocinnimate, cadaverine, and putrescine are markers of periodontal disease that were reduced in saliva when supragingival plaque was removed, suggesting they were largely produced by bacteria (Sakanaka et al. 2017), and we detected each of these molecules. This demonstrates that bacterial metabolic products are present in calculus, and may offer insight into mature biofilm activity. This could be useful in studying how bacterial metabolism influences oral disease, as periodontal disease-associated oral plaque has community structure and activity much more similar to that of fully mature biofilms such as are found in calculus, than to healthy site subgingival plaque or supragingival plaque (Wade 2013), yet the presence of calculus alone is not a reflection of periodontal disease status (i.e., three of the five modern calculus samples were collected from teeth with no evidence of periodontal disease).

In sum, our results demonstrate that dental calculus contains an abundance of endogenous and exogenous metabolites, and that a wide range of these metabolites preserve well through time. Dental calculus therefore has significant potential to provide novel insights into human diet, physiology, and microbiome activity in both modern and historic samples, permitting human evolutionary and human-microbiome co-evolutionary studies with a deep-time perspective. Larger sample sizes and samples from additional temporal and cultural contexts as well as from varying burial and storage conditions are needed to further address metabolite preservation and presence/absence patterns. The excellent preservation of dental calculus in archaeological collections, however, means that there is ample opportunity to expand metabolite-based studies of dental calculus into the recent and distant past.

## Acknowledgments

The authors thank Mark Gibson, Tom Gilbert, Chris Gosden, Stuart Gould, Lauren McIntyre, Anita Radini, William Wade, and Helen Webb for assistance with sample and data collection. We would like to acknowledge The Danish National High-Throughput DNA Sequencing Centre for sequencing the samples. This work was supported by the Oxford University Fell Fund 143/108 (to G.L. and C.W.), the European Research Council ERC-2013-StG 337574- UNDEAD (to G.L.), the National Institutes of Health R01GM089886 (to C.M.L., C.W., and K.S.), the National Science Foundation BCS-1516633 and BCS-1643318 (to C.W.), and R35GM118110 (to J.J.C.) and the National Library of Medicine T15LM007359 (to K.A.O.).

## Data availability

All data generated or analyzed during this study are included in this published article (and its supplementary information files).

## Supplementary Tables

**Table S1**. Raw UPLC-MS/MS data from Metabolon. Tab 1 (OrigScale) contains data normalized in terms of raw area counts. Tab 2 (MassNormImp) contains data normalized as follows: values for each sample are normalized by sample mass available/utilized for extraction, then each biochemical is rescaled to set the median equal to 1, and missing values are imputed with the minimum. (Excel sheet).

**Table S2**. Raw Data from the University of Wisocnson-Madison. Tab 1 Metabolites detected by GC-MS. Tab 2 Metabolites detected by UPLC-MS/MS. (Excel sheet).

**Table S3.** Differential abundance of UPLC-MS/MS-identified metabolites between modern and historic dental calculus. (Word doc).

## Supplementary Figures

**Fig. S1.**
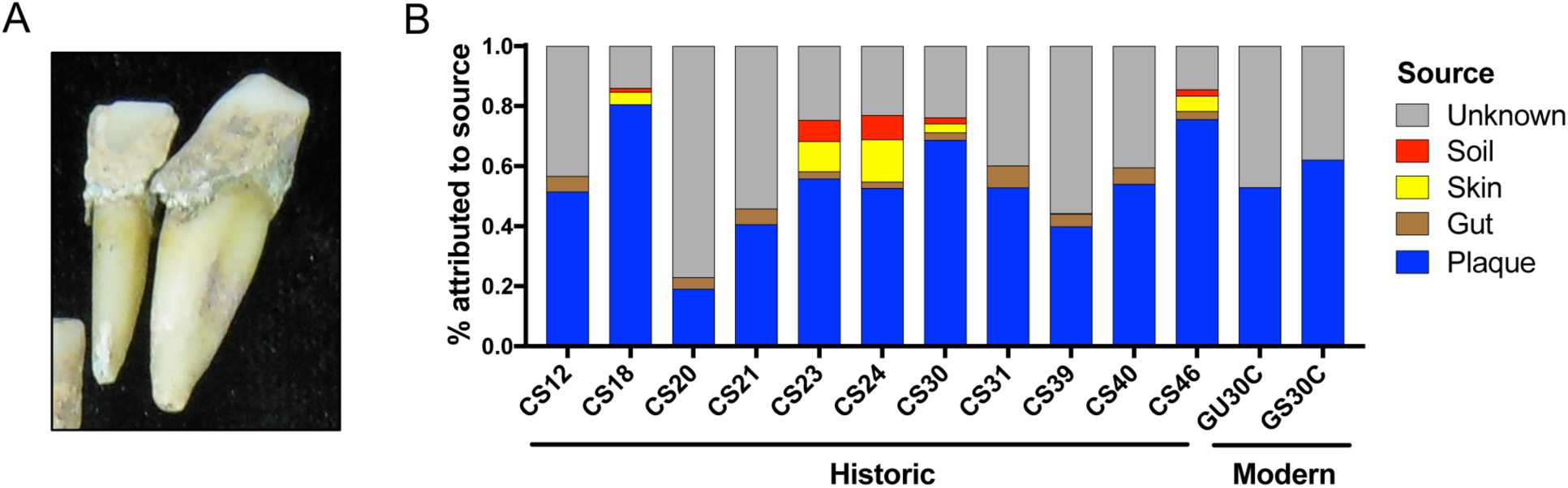
Historic calculus samples contain oral bacterial community profiles. **a** Historic calculus on teeth from CS18 prior to sampling. **b** Percent of bacterial community in calculus samples attributable to distinct environmental sources by SourceTracker analysis. Modern samples from Ziesemer et al. (2015), demonstrate that a high proportion of the microbial community assigned to an “Unknown” source is characteristic of dental calculus. CS6 failed to build DNA libraries with the AccuPrimePFX polymerase.

**Fig. S2.**
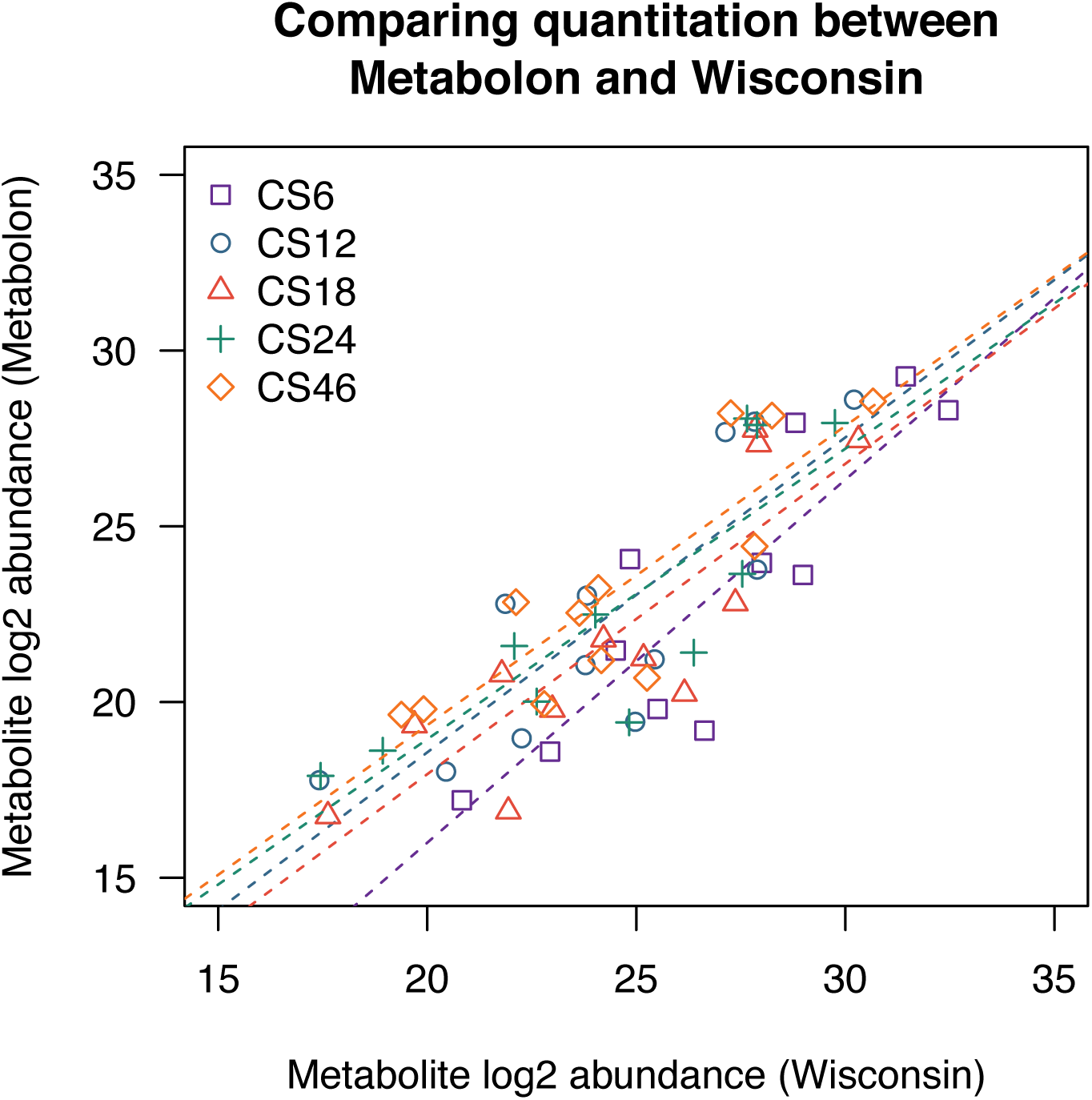
Comparison of quantified metabolites from samples analyzed by Metabolon, Inc. and Wisconsin-Madison. Five calculus samples were analyzed by GC-MS and LC-MS in Madison, Wisconsin; the relative quantitation obtained on these methods was significantly correlated with results from Metabolon, Inc. (linear regression, p < 0.001).

**Fig. S3.**
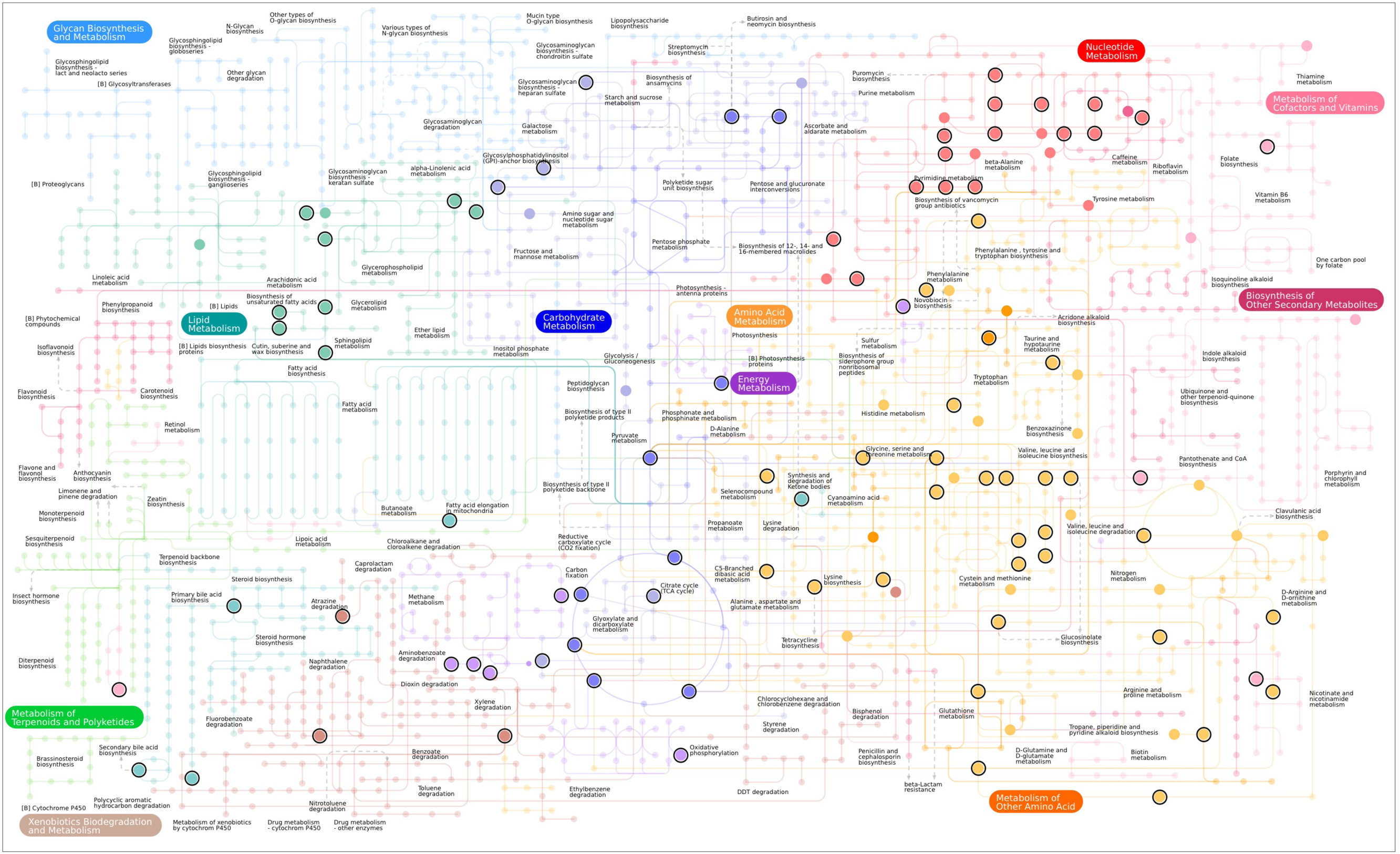
Representation of diverse metabolic pathways preserved in calculus. Large circles represent metabolites identified in calculus. Black rings around large circles indicate the metabolite was detected in historic calculus.

**Fig. S4.**
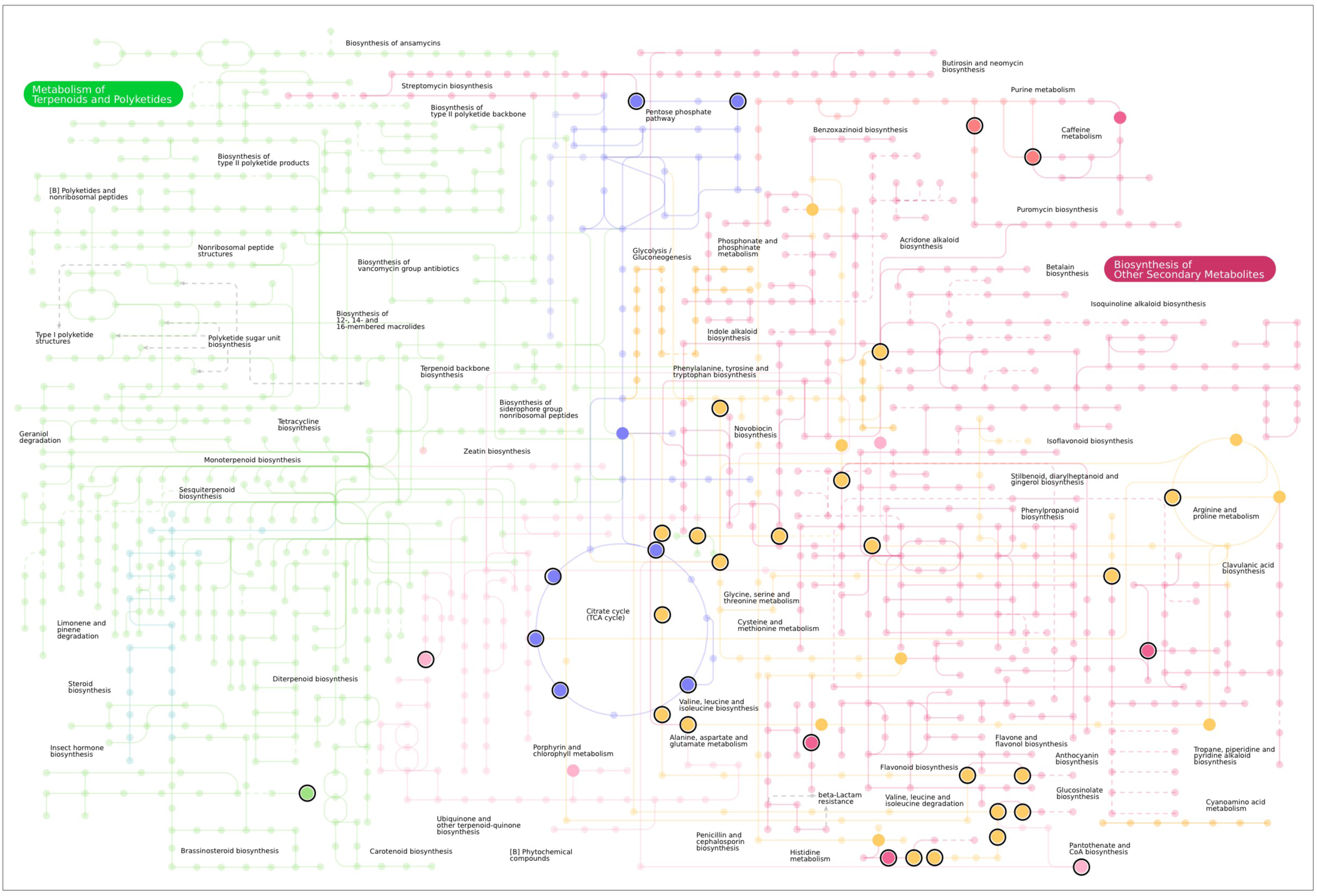
Representation of pathways for biosynthesis of secondary metabolites preserved in calculus. Large circles represent metabolites identified in calculus. Black rings around large circles indicate the metabolite was detected in historic calculus.

**Fig. S5.**
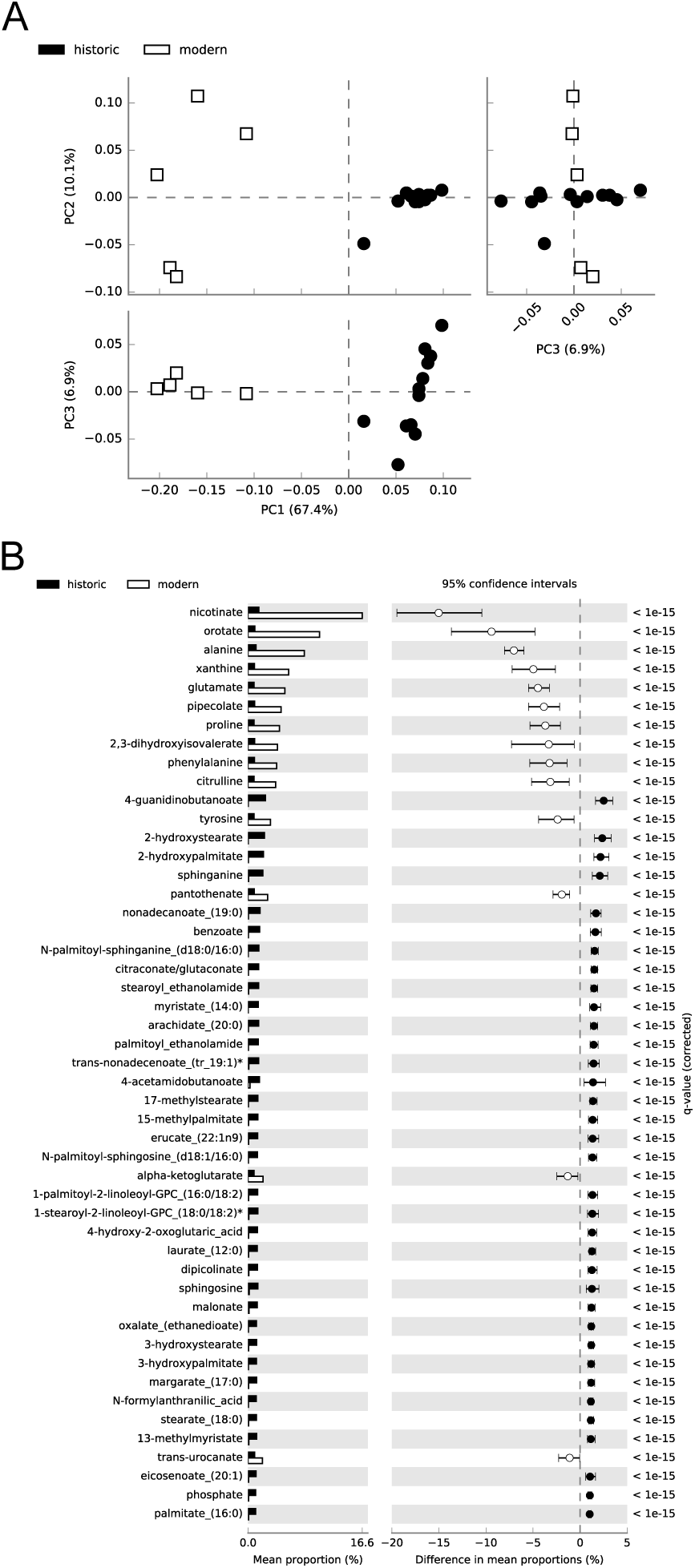
Differences exist in mean proportions of metabolites universally detected in all historic and modern dental calculus samples. **a** Principal components analysis distinctly separates modern and historic calculus samples. **b** Metabolites with significant differences (q ≤ 0.05, effect size of ≥1.0) in mean proportions between historic and modern calculus.

**Fig. S6.**
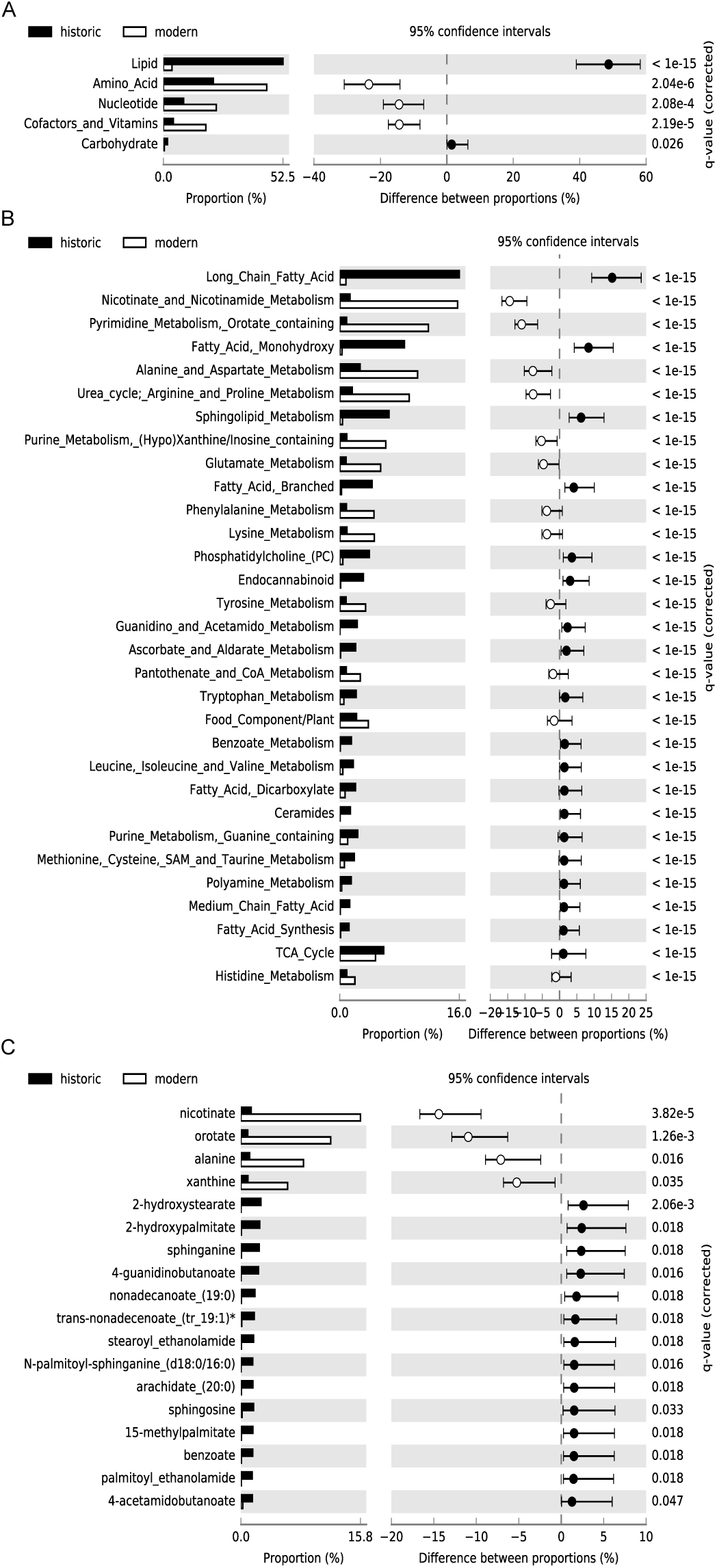
Differences exist in proportions of super-pathways, sub-pathways, and metabolites universally detected in all historic and modern dental calculus samples. Significantly different (q ≤ 0.05, effect size of ≥1.0) proportions of **a** Super-pathways, **b** Sub-pathways, and **c** Individual metabolites between historic and modern samples.

**Fig. S7.**
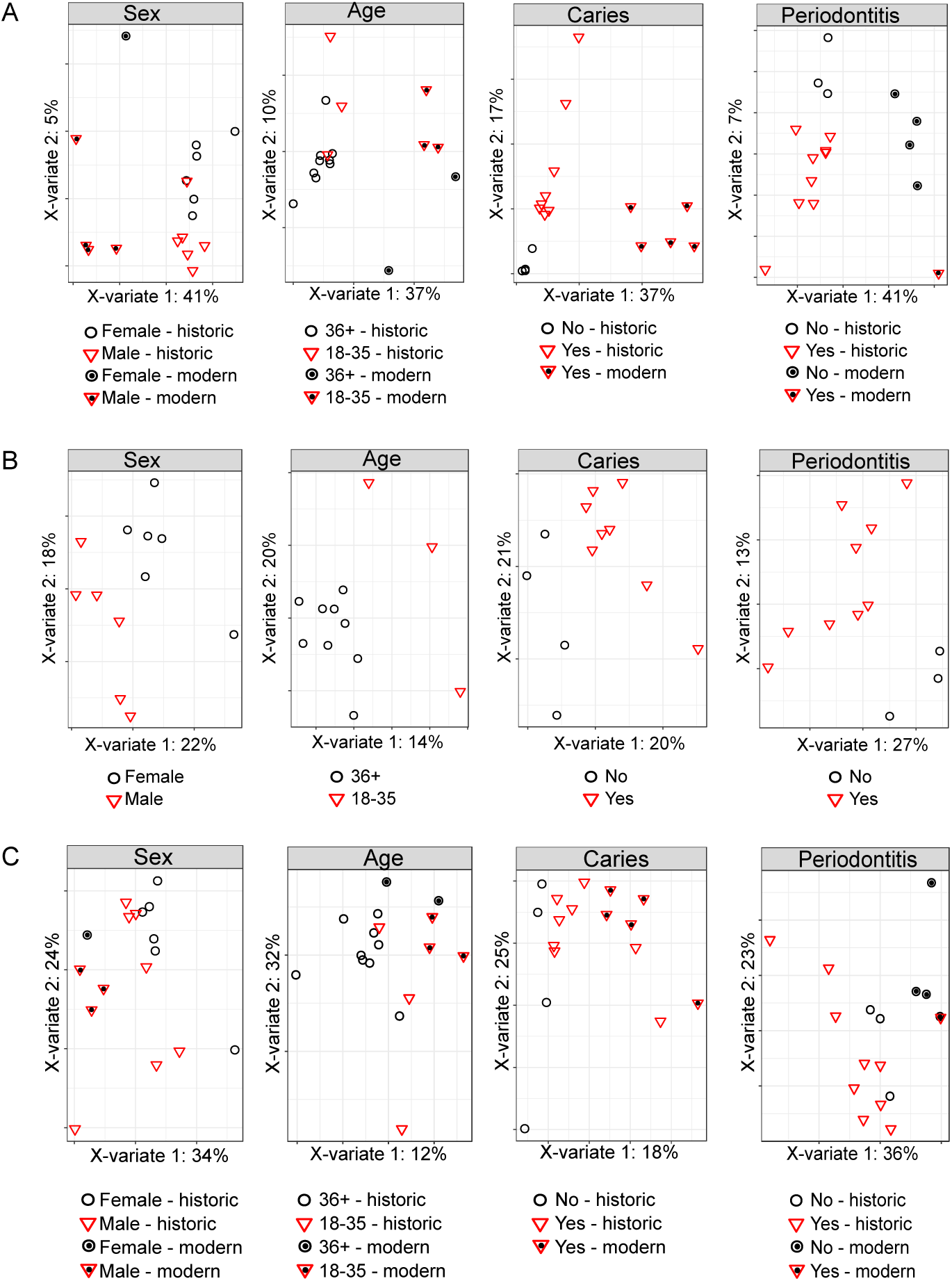
Partial least squares discriminant analysis of metabolites universally present in calculus samples. **a** Calculus samples cluster based on time period rather than biological category (sex, age, caries status and periodontal disease status) when including metabolites detected in all seventeen calculus samples. **b** Historic calculus samples cluster based on biological category (sex, age, caries status and periodontal disease status) when including metabolites detected in all twelve historic samples. **c** When using only universally detected metabolites in the *Lipid* and *Energy* categories (pathways with the best representation in historic samples), it was still not possible to discriminate samples based on biological category, although the separation between historic and modern samples was reduced slightly compared to **a.**

**Fig. S8.**
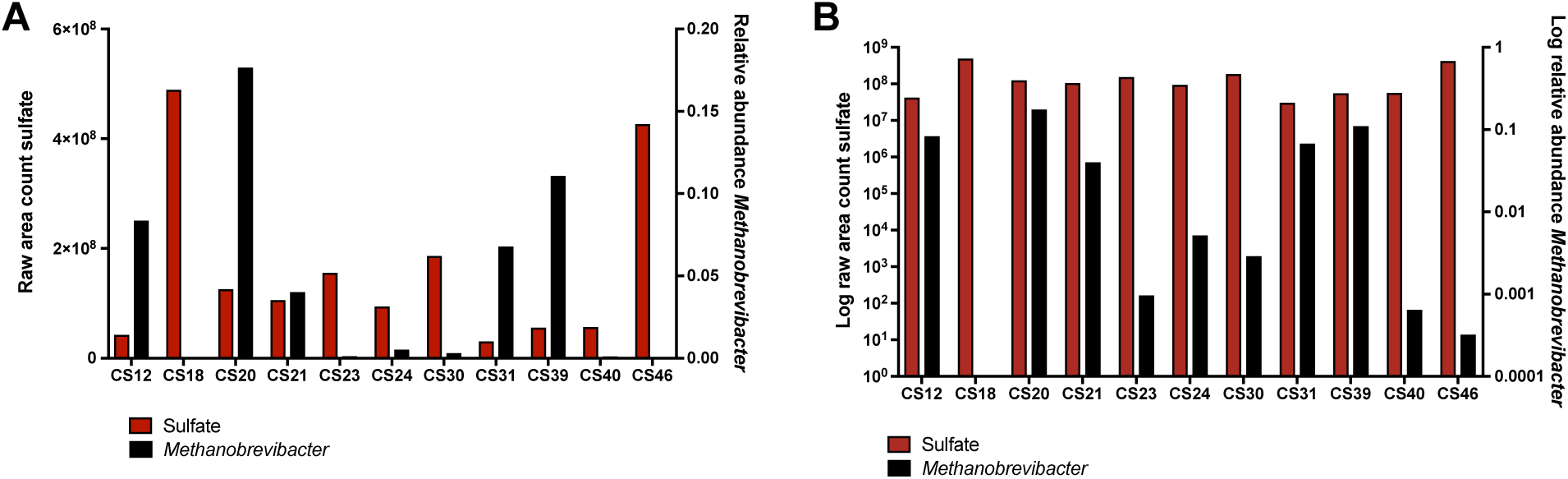
Comparison of sulfate abundance and the relative abundance of the oral sulfate-producing genus, *Methanobrevibacter*, shown using **a** linear and **b** log scales. *Methanobrevibacter* relative abundance was estimated using 16S rRNA gene sequence counts, and no correlation was observed with sulfate abundance. *Methanobrevibacter* was not detected in CS18.

## Supplementary Materials

### 1. Supplementary Figures

### 2 Supplementary Materials and Methods

#### 2.1 Calculus collection and preparation

All of the skeletons were from earth cut graves and had either been contained within wooden coffins, subsequently decomposed, or had been buried in shrouds uncoffined. The surfaces of the teeth were cleaned with 5% NaOCl followed by water prior to sampling to remove traces of burial soil, and sampling was performed wearing gloves and a mask over the nose and mouth. Calculus samples were collected in individual tubes on site, and removed to the Research Laboratory for Archaeology and the History of Art at the University of Oxford where ∼20 mg was subsampled, placed in a new tube and crushed by micropestle. Crushed historic calculus samples were sent for metabolomics analyses without further processing.

#### 2.2 Genetic Authentication of a Preserved Oral Microbiome in Historic Samples

Shotgun Illumina libraries were constructed following previously described methods (Meyer and Kircher 2010) with AccuPrime PFX polymerase (Invitrogen), and sequenced on an Illumina HiSeq2500 at the University of Copenhagen National High-Throughput DNA Sequencing Centre. Prior to analysis, reads were de-mulitplexed, quality-checked, and trimmed of adapters using AdapterRemoval v.1 (Lindgreen 2012) with the following non-default parameters: --maxns 0, --trimns, --trimqualities --minquality 30, --minlength 25, --collapse, and --minalignmentlength 10. To identify 16S rRNA gene reads in the metagenomic dataset, reads were aligned to the Greengenes v. 13.8 database using bowtie2 (Langmead and Salzberg 2012).

#### 2.3 Sample Preparation for Mass Spectrometry at Metabolon, Inc.

Samples (∼20 mg) were decalcified in 0.5M EDTA, centrifuged to pellet debris, and supernatant prepared using the automated MicroLab STAR® system from Hamilton Company. Several recovery standards were added prior to the first step in the extraction process for QC purposes. To remove protein, dissociate small molecules bound to protein or trapped in the precipitated protein matrix, and to recover chemically diverse metabolites, proteins were precipitated with methanol under vigorous shaking for 2 min (Glen Mills GenoGrinder 2000) followed by centrifugation. The resulting extract was divided into five fractions: two for analysis by two separate reverse phase (RP)/UPLC-MS/MS methods with positive ion mode electrospray ionization (ESI), one for analysis by RP/UPLC-MS/MS with negative ion mode ESI, one for analysis by HILIC/UPLC-MS/MS with negative ion mode ESI, and one sample was reserved for backup. Samples were placed briefly on a TurboVap® (Zymark) to remove the organic solvent. The sample extracts were stored overnight under nitrogen before preparation for analysis.

#### 2.4 QA/QC at Metabolon, Inc.

Several types of controls were analyzed in concert with the experimental samples: a pooled matrix sample generated by taking a small volume of each experimental sample (or alternatively, use of a pool of well-characterized human plasma) served as a technical replicate throughout the data set; extracted water samples served as process blanks; and a cocktail of QC standards that were carefully chosen not to interfere with the measurement of endogenous compounds were spiked into every analyzed sample, allowed instrument performance monitoring and aided chromatographic alignment. Instrument variability was determined by calculating the median relative standard deviation (RSD) for the standards that were added to each sample prior to injection into the mass spectrometers. Overall process variability was determined by calculating the median RSD for all sample metabolites (i.e., non-instrument standards) present in 100% of the pooled matrix samples. Experimental samples were randomized across the platform run with QC samples spaced evenly among the injections.

#### 2.5 Ultrahigh Performance Liquid Chromatography-Tandem Mass Spectroscopy (UPLC-MS/MS) at Metabolon, Inc.

All methods utilized a Waters ACQUITY ultra-performance liquid chromatography (UPLC) and a Thermo Scientific Q-Exactive high resolution/accurate mass spectrometer interfaced with a heated electrospray ionization (HESI-II) source and Orbitrap mass analyzer operated at 35,000 mass resolution. The sample extract was dried then reconstituted in solvents compatible to each of the four methods. Each reconstitution solvent contained a series of standards at fixed concentrations to ensure injection and chromatographic consistency. Two aliquots were analyzed using acidic positive ion conditions, the first chromatographically optimized for more hydrophilic compounds and the second chromatographically optimized for more hydrophobic compounds. In the first method the extract was gradient eluted from a C18 column (Waters UPLC BEH C18- 2.1x100 mm, 1.7 µm) using water and methanol, containing 0.05% perfluoropentanoic acid (PFPA) and 0.1% formic acid (FA). In the second method, the extract was gradient eluted from a C18 column using methanol, acetonitrile, water, 0.05% PFPA and 0.01% FA and was operated at an overall higher organic content. Another aliquot was analyzed using basic negative ion optimized conditions using a separate dedicated C18 column, and were gradient eluted from the column using methanol and water with 6.5mM Ammonium Bicarbonate at pH 8. The fourth aliquot was analyzed via negative ionization following elution from a HILIC column (Waters UPLC BEH Amide 2.1x150 mm, 1.7 µm) using a gradient of water and acetonitrile with 10mM Ammonium Formate, pH 10.8. The MS analysis alternated between MS and data-dependent MS^n^ scans using dynamic exclusion. The scan range varied slighted between methods but covered 70-1000 m/z.

#### 2.6 Data Extraction, Compound Identification, Quantification, and Normalization at Metabolon, Inc.

Raw data was extracted, peak-identified and QC processed using Metabolon’s hardware and software. Compounds were identified by comparison to library entries of purified standards or recurrent unknown entities. The MS/MS scores are based on a comparison of the ions present in the experimental spectrum to the ions present in the library spectrum. Library matches for each compound were checked for each sample and corrected if necessary. Peaks were quantified using area-under-the-curve. Each compound was corrected in run-day blocks by registering the medians to equal one (1.00) and normalizing each data point proportionately.

#### 2.7 Further characterization of historic calculus by GC-MS and UPLC-MS/MS

For GC-MS analysis, dried extract was derivatized for 90 min with 20 mg/mL methoxyamine hydrochloride in pyridine at 20°C (10 uL) and then with MSTFA for 30 min at 37°C (10 uL). Samples were analyzed by GC-Orbitrap; 1 uL of sample, split 1:10, was injected onto a TraceGOLD TG-5SilMS GC column (cat. no. 26096-1420, Thermo Scientific). Temperature was held at 50°C for 1 min, then ramped to 320°C at rate of11°C/min, then held at 320°C for 4.40 min. Molecules were analyzed with positive electron-impact (EI)-Orbitrap full scan of 50-650 m/z range. Raw files were analyzed using Thermo Scientific’s Tracefinder 4.0 deconvolution plugin and unknown screening quantification tool. Deconvolved peaks were searched against NIST 2014 and in-house high-resolution GC libraries; retention index was used to filter search hits. A single quant-ion was used for quantification of deconvolved peaks. Quantified peaks in samples were included if they were at least 10-fold greater than peaks quantified in solvent blanks. For lipid analysis, dried extract was resuspended in 65:30:5 isopropanol:acetonitrile:water. Lipid LC-MS analysis was performed on a Water’s Acquity UPLC CSH C18 Column (2.1 mm x 100 mm) with a 5 mm VanGuard Pre-Column Mobile coupled to a Q Exactive Focus. Mobile phase A consisted of 70% acetonitrile and 30% water with 10 mM Ammonium acetate and 0.025% acetic acid. Mobile phase B consisted of 90% isopropanol and 10% acetonitrile with 10 mM ammonium acetate and 0.025% acetic acid. Samples were separated using the following 30 min gradient: 2% B for 2 min (0.4 mL/min), increased to 30% B over next 3 min (0.4 mL/min), increased to 50% B over next 1 min (0.4 mL/min), increased to 85% B over next 14 min (0.4 mL/min), increased to 99% B over next 1 min and held at 99% B for 7 min (0.3 mL/min); then returned to 2% to equilibrate for 2 min (0.4 mL/min). Samples were analyzed on the previous gradient using in first positive ion mode and then negative ion mode electrospray ionization (ESI) with full scan MS1 (150-1600 Th) collected 17,000 resolving power (at 400 m/z) for 0-30 min and top-2 data dependent MS2 scans fragmented with stepped normalized collision energy (20-40%). Raw files were quantified using the Thermo Compound Discoverer^TM^ 2.0 application with peak detection, retention time alignment, and gap filling. Only peaks 10- fold greater than solvent blanks were included in the later analysis. Identification was aided by in-house software and lipid libraries; only compounds with definitive MS/MS evidence were assigned with identification.

